# Ecological insights and management implications of the global migratory connectivity of green turtles

**DOI:** 10.1101/2025.09.07.674769

**Authors:** Jaime Restrepo, Dina Nisthar, Harris Wei-Khang Heng, Lily K Bentley, Corrie Curtice, Sarah DeLand, Ei Fujioka, Patrick N. Halpin, Sarah K. Poulin, Anthony J Richardson, Jeffrey A. Seminoff, Roldna A. Valverde, Daniel C. Dunn

## Abstract

**Aim:** Green turtles are a widely distributed and highly migratory species, despite extensive data on the movement of green turtles, there is no global synthesis on the subject, limiting a holistic understanding of their movement. Based on three decades of published literature, we present the first global model of migratory connectivity for green turtles.

**Location:** Global.

**Time Period:** 1990–2022.

**Major Taxa Studied:** Green Sea Turtle (*Chelonia mydas*)

**Methods:** We conducted a structured literature review extracting georeferenced information on the movement of green turtles from 1990 to 2022, aggregating this information into a single connectivity model, defining sites and “metasites” as nodes of connectivity. We then evaluate the connectivity routes from nesting areas to foraging sites for each RMU, identifying those trajectories moving outside and across the boundaries of these areas.

**Results:** We found an increasing number of studies assessing movement of green turtles, with a total of 113 sources of migratory connectivity information. We identified 474 sites, representing locations where green turtles were observed (124 of these being nesting sites). Migratory connections from nesting sites ranged from resident turtles never leaving the area, to rookeries linked to 13 different sites, some over 5,000 km apart. This long-distance connectivity exposes populations to threats across disparate locations. Most connections traversed national jurisdictions, including crossing different Regional Management Units

**Main Conclusions:** We compiled the largest available dataset describing movement of green turtles worldwide and present the first global model of their migratory connectivity. This model provides ecological insights into regional differences in life histories, identifies geographic and demographic gaps in sampling, and provides baseline information on connectivity to support transboundary management of green turtle populations. The study highlights the need for larger collaborative efforts to aggregate knowledge beyond local jurisdictions, to inform and align effective management measures to protect this threatened species.

## Introduction

Connectivity is a fundamental component of spatial ecology and can inform conservation (Virtanen et al. 2020). In both landscape and metapopulation ecology, connectivity is used to describe the degree of movement or exchange of organisms between habitats (Dibacco et al. 2006). Animal movement between habitats is essential in the evolution of migratory species, as it influences accessibility of resources, which controls how a species uses different habitats throughout its life cycle. Habitat connectivity has a profound influence at a population level, affecting genetic structure, migratory movements, and the response to external stressors such as climate change (Baguette & Van Dyck 2007; Kool et al. 2013).

Limited movement amongst diverse habitats, may affect isolated ecosystems, reducing their ability to support animal populations.

There are various types of ecological spatial connectivity acting at multiple spatial scales (Bryan-Brown et al. 2017). One of the most important is migratory connectivity, which describes how animal movement creates geographic links across seasons as individuals roam long distances to congregate, breed and feed (Webster et al. 2002). This concept has important consequences for the ecology, evolution and conservation of migratory species, as successful reproduction depends on key habitats, which they occupy seasonally throughout their life. Thus, identifying breeding and non-breeding habitats, and connections between them, is critical for the implementation of effective conservation and management strategies (Dunn et al. 2019).

Assessing migratory connectivity for marine megavertebrates is demanding, given the difficulties associated with tracking elusive animals moving over vast, opaque, three-dimensional geographical ranges (Vander Zanden et al. 2015; Jeffers & Godley 2016; Horton et al. 2017). Over the past few decades, advances in remote monitoring technology have greatly improved our ability to study animal movement, allowing researchers to track the cryptic dispersive stages of marine animals (Hussey et al. 2015; Bryan-Brown et al. 2017). The rapid development of satellite telemetry technology has created unprecedented opportunities to monitor animal movements remotely, providing previously inaccessible information on area use, migratory strategies, non-breeding areas and connectivity (Hazen et al. 2012; Vander Zanden et al. 2015; Horton et al. 2017). This is particularly true for animals that are large (so can easily carry satellite tags), easy to capture, and long-lived, such as marine turtles (Wallace et al. 2010).

Green turtles (*Chelonia mydas)* are widely distributed and undertake long migrations that regularly cross multiple national jurisdictions (Seminoff et al. 2015). The transboundary nature of their life histories complicates conservation strategies, as it can be difficult to align management across jurisdictions (Roberson et al. 2021) and governance of the high seas remains fragmented (Ban et al. 2014). To address these challenges, in 2010 the IUCN Marine Turtle Specialist Group developed the concept of marine turtle Regional Management Units (RMUs). RMUs provide a globally consistent management framework that defines turtle assemblages below the level of species but above the level of breeding rookeries (Wallace et al. 2010, 2023). Within RMUs, green turtles have long-term fidelity to foraging sites (Metz et al. 2020; Pilcher et al. 2020; Siegwalt et al. 2020; Webster et al. 2022) and nesting areas (Lee et al. 2007; Esteban et al. 2017; Shimada et al. 2021). Although many studies assess the movement of green turtles (e.g., Godley et al. 2008; Shaver et al. 2013), gaps remain in our understanding of migratory connectivity for most populations in the world (Godley et al. 2010).

Here, we synthesise global scale data, collected on green turtle movements over three decades to identify migratory connectivity linking key nesting and foraging areas. By synthesising and integrating this information, we derive the first global model of the currently known migratory connectivity for green turtles. We also quantify global biases in connectivity data for this species, discerning between different age and sex classes. The model provides novel ecological insights into regional differences in life histories, identifies geographic and demographic gaps in sampling, and provides baseline information on connectivity to support transboundary management of green turtle populations worldwide.

## Methods

### Literature search and data extraction

We conducted a structured literature review using the Web of Science and Scopus databases to search for literature on ecological connectivity data on the movement of green turtles, following Kot et al. (2023). Our search included peer-reviewed articles, book chapters, scientific reports, and conference proceedings from 1990–2022. Prior to 1990, there was very limited satellite tracking of marine species. Within the Web of Science database, all searches were sorted by topic to identify every publication in English over the 33-year period of study within the subscribed databases (WOS, BCI, CCC, DRCI, DIIDW, KJD, MEDLINE, RSCI, SCIELO, ZOOREC). Similarly, within the Scopus database, all searches were conducted by keyword, title, and subject area. The search string combined species name with tagging or sampling techniques and a variation of terms indicating movement or connectivity.

"<*Chelonia mydas*> AND <GREEN turtle> AND (telemetry OR tag* OR isotop* OR genetic* OR mark OR recapture) AND (migrat* OR connect* OR move* OR feed* OR forag* OR breed* OR dispers* OR nest* OR aggregat* OR ground* OR site* OR corridor OR route* OR track* OR winter* OR habitat*)" This search string captured a wide range of information, including different life stages and various connectivity sampling methods. After standardising the methods to extract information from relevant references, nine reviewers collected detailed information for each article including: 1) latitude and longitude of deployment and the final destination of each tagged turtle; 2) scale of movement around each deployment location and final destination; 3) number of turtles monitored, including their sex (i.e. female, male or undetermined) and life stage (i.e. adult, adult breeder, sub-adult, juvenile or unknown); 4) behaviour at deployment site and final destination (i.e. nesting, foraging, or migrating); and 5) method used to assess movement and connectivity (e.g., mark-recapture, satellite telemetry, acoustic telemetry). We extracted site and connectivity information, where sites were defined as georeferenced areas used by animals, typically associated with a specific behaviour or activity (e.g., foraging, breeding, nesting). Connections were defined as the links among these sites (adapted from Kot et al. 2023).

### Global model of migratory connectivity

Given the variety of research approaches implemented over time and the lack of detailed information describing animal movements in some studies, we followed Bentley et al. (2025) to identify and aggregate georeferenced sites from the literature where green turtle movements were reported. Sites were delineated as a circle with a 1°-, 5°- or 10°-radius, where a specific behaviour had been identified or where undefined “observations” of green turtles had been detected. Aggregation of sites across studies was necessary, as many studies depict the same site. To pool information from multiple studies into a single connectivity model, we aggregated sites into “metasites” based on four objectives: 1) aggregate information from nearby locations used by green turtles; 2) to preserve information on large-scale (>500 km) connections revealed by animal movements; 3) maintain spatial resolution of information on critical reproductive life history stages; and 4) provide a minimum estimate of animals recorded within a site and moving between sites.

To address these objectives, we implemented a five-step process for site aggregation (full details reported in Bentley et al. [2025]). In summary, we iteratively evaluated all sites by selecting a “lookup site” and subsequently identifying sites that could be aggregated with it whose centroid fell within the buffer of the lookup site. To ensure that regional connectivity information was not lost in the aggregation process, we prevented sites associated with connections that described movement across distances >500 km from being aggregated together. Subsequently, to develop a metasite, we implemented an iterative subprocess in which we calculated the new metasite centroid as the mean latitude and longitude from those aggregated sites. Any sites included in the new metasite were no longer considered for aggregation into subsequent metasites.

Given that many tracking studies repurposed or expanded existing datasets, we identified potentially duplicated data to ensure that our sample size estimates were conservative and representative of a minimum estimate of known connectivity. We considered duplicate sites those that: 1) had identical latitudes and longitudes and were from the same study; 2) sites that were aggregated together in a metasite and had a connection between them, so as not to double count animals moving within a metasite; and 3) sites extracted from references with at least one author with the same surname where the temporal scope of the data in one study was a subset of that in the other study (see Bentley et al. 2025 for figures describing the workflow).

### Migratory connectivity and Regional Management Units

Regional management Units can overlap in certain areas, which may translate into conservation gaps for the status of different populations (Wallace et al. 2025). Based on the geographical location for each site and metasite, we assigned them to specific RMUs for green turtles across the world (Wallace et al. 2023). We then evaluated the connectivity between nesting and foraging sites for each RMU, identifying those trajectories moving outside and across the boundaries of these areas.

## Results

### Literature search and data extraction

We synthesised information on global movement, behaviour, and migratory connectivity of green turtles from 113 different sources. These studies were within the known historical distribution of the green turtles in the Atlantic (42.5% of references, n = 48), Indian (17.7%, n = 20) and Pacific (39.8%, n = 45) oceans. The annual number of publications increased over the 33-year search period, with nearly half (47.8%) of all references published in the last decade (Figure 1). A total of 7,360 green turtles were tagged in studies to assess their movement, with flipper tags making up the vast majority (83.6%, n = 6155), while 998 turtles (13.6%) were equipped with satellite telemetry tags. The remaining monitored turtles were tagged using data loggers, acoustic tags, or a combination of these techniques. Most tagged animals were female (85.5%), predominantly sampled at nesting beaches. Tracking devices tended to be applied individually, although in a few cases (<2.0%) studies reported a combination of different tag types (e.g., acoustic and satellite tags) as an experimental treatment to assess the accuracy of the instruments (Table S1).

**Figure 1.**
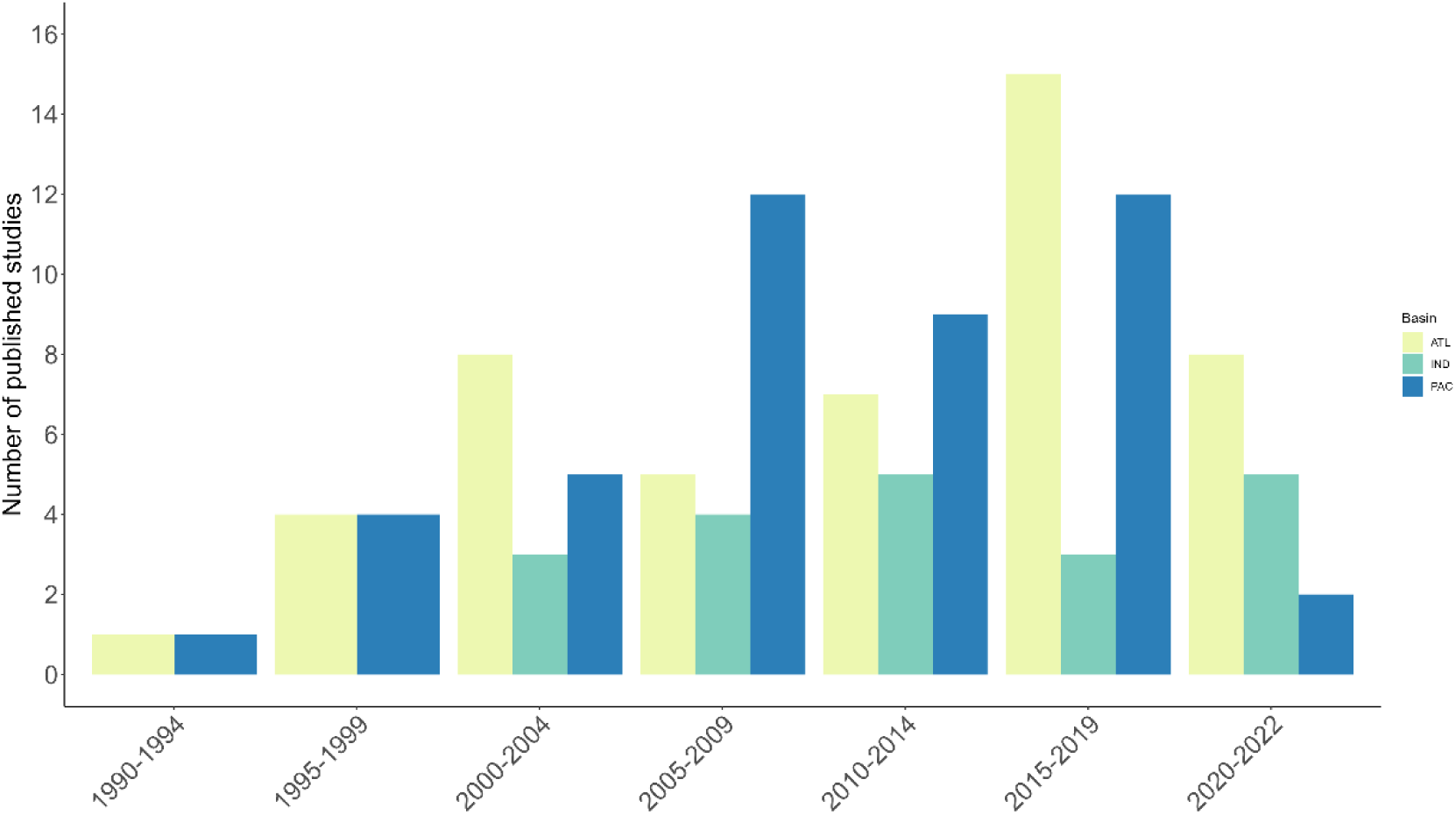
Number of published studies assessing green turtle (*Chelonia mydas*) movement separated by oceans (ATL = Atlantic; IND = Indian; PAC = Pacific) for all five-year intervals between 1990 and 2022.

Various questions related to green turtle movement ecology have been addressed during the past three decades. More than a third of studies (37.2%, n = 42) examined the post-nesting migrations of female green turtles after departing known rookeries. Over a quarter of studies (27.4%, n = 31) explored swimming behaviour and migratory strategies, primarily examining the dispersal of younger turtles or evaluating the behaviour and movement of turtles reared in captivity. Other areas of study included, assessment of habitat use (17.6%, n = 20), foraging home range (12.4%, n = 14), experimental trials (3.6%, n = 4) and interaction with fisheries (1.8%, n = 2; Table S1).

Most studies tracked female breeding turtles using telemetry devices (70.1%). Several studies tracked juveniles of undetermined sex (18.6%), and turtles of mixed age classes (10.4%) -based on the curved length of their carapace and tail size- (Figure 2a; 2b). Only few studies (0.9%) conducted in-water, tracked male turtles from their foraging habitats; information on these groups still represent a major gap in knowledge for green turtles’ movement. Information extracted from these studies predominantly included adults (75.7%), most of them undertaken at breeding sites (72.4%). The remaining studies studied subadult (12.5%), juvenile (8.9%) and turtles of unknown age class (2.9%) tracked at sea after being released in foraging grounds (Figure 2b).

**Figure 2.**
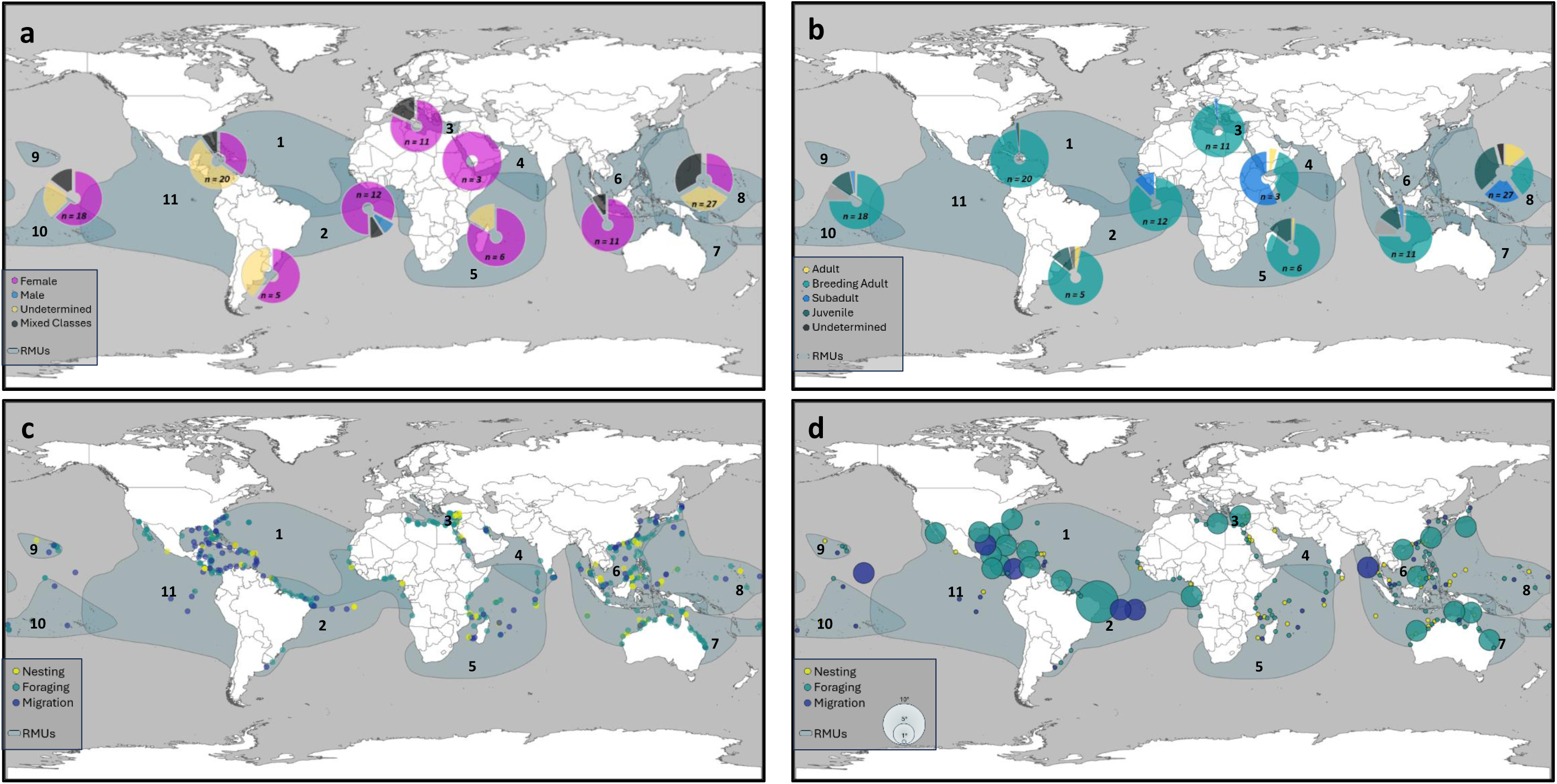
Green turtle (*Chelonia mydas*) information extracted from the literature, showing proportion of references from geographical regions. a) Proportion of turtles studied in terms of their sex: Female, Male, Undetermined, Mixed Classes. b) Proportion of turtles studied in different regions based on life-stage: Adult, Adult Breeding, Subadult, Juvenile, Undetermined). c) Sites where turtles were tagged and released or observed after tagging, subdivided by behaviour: Nesting, Foraging, Migrating. d) Metasites of green turtles after aggregation of information subdivided by behaviour: Nesting, Foraging, Migrating. Blue shaded areas represent Regional Management Units (RMUs) for green turtles; 1: North Atlantic, 2: South Atlantic, 3: Mediterranean, 4: Northwest Indian, 5: Southwest Indian, 6: East Indian and Southeast Asia, 7: Southwest Pacific, 8: West Central Pacific, 9: North Central Pacific, 10: South Central Pacific, 11: East Pacific. (Wallace et al. 2023).

### Global model of migratory connectivity

We identified 474 sites, representing both locations where green turtles were initially tagged and locations recorded after migration. Of these locations, 124 sites were breeding or nesting sites, and 186 were foraging sites. Few locations were detected for behaviours such as migration, stopover, or wintering. A total of 87 sites had a combination of two or more different behaviours (Figure 2c).

After aggregating overlapping and nearby sites, we identified 189 unique metasites. From these, 55 reproductive grounds were preserved as 1° metasites, whilst most foraging areas were aggregated in 5° metasites. Brazil has the only 10° metasite, aggregating information from 18 different foraging grounds off the coastal state of Rio Grande Do Norte, constituting the largest foraging metasite identified of this network. We aggregated information from several studies tracking green turtles to this area from distant rookeries, establishing connections between the metasite and various nesting locations across the Atlantic Ocean. Nesting females tracked from Guinea Bissau in West Africa (∼2,800 km), Ascension Island (∼2,400 km) and Costa Rica (>5,000 km) gathered at the Brazilian coast to feed (Figure 2d).

Globally, we identified 206 connections between metasites, primarily showing movements from nesting sites towards foraging areas (Figure 3a). These connections included relatively small-scale (∼400 km) movements of individuals in the Red Sea and Caribbean Sea (noting that any movements <100 km would be aggregated within a given site). There were also larger-scale movements (∼3,000 km) of individuals from Paraiba in Brazil to Guinea on the west coast of Africa, and from Lacepede Island in Western Australia to the Gulf of Carpentaria between northern Australia and Papua in Indonesia. Further, the longest recorded migration belonged to green turtles traveling >5,000 km from nesting beaches in Micronesia towards foraging habitats off the coast of peninsular Malaysia (Figure 3a, 4c).

**Figure 3.**
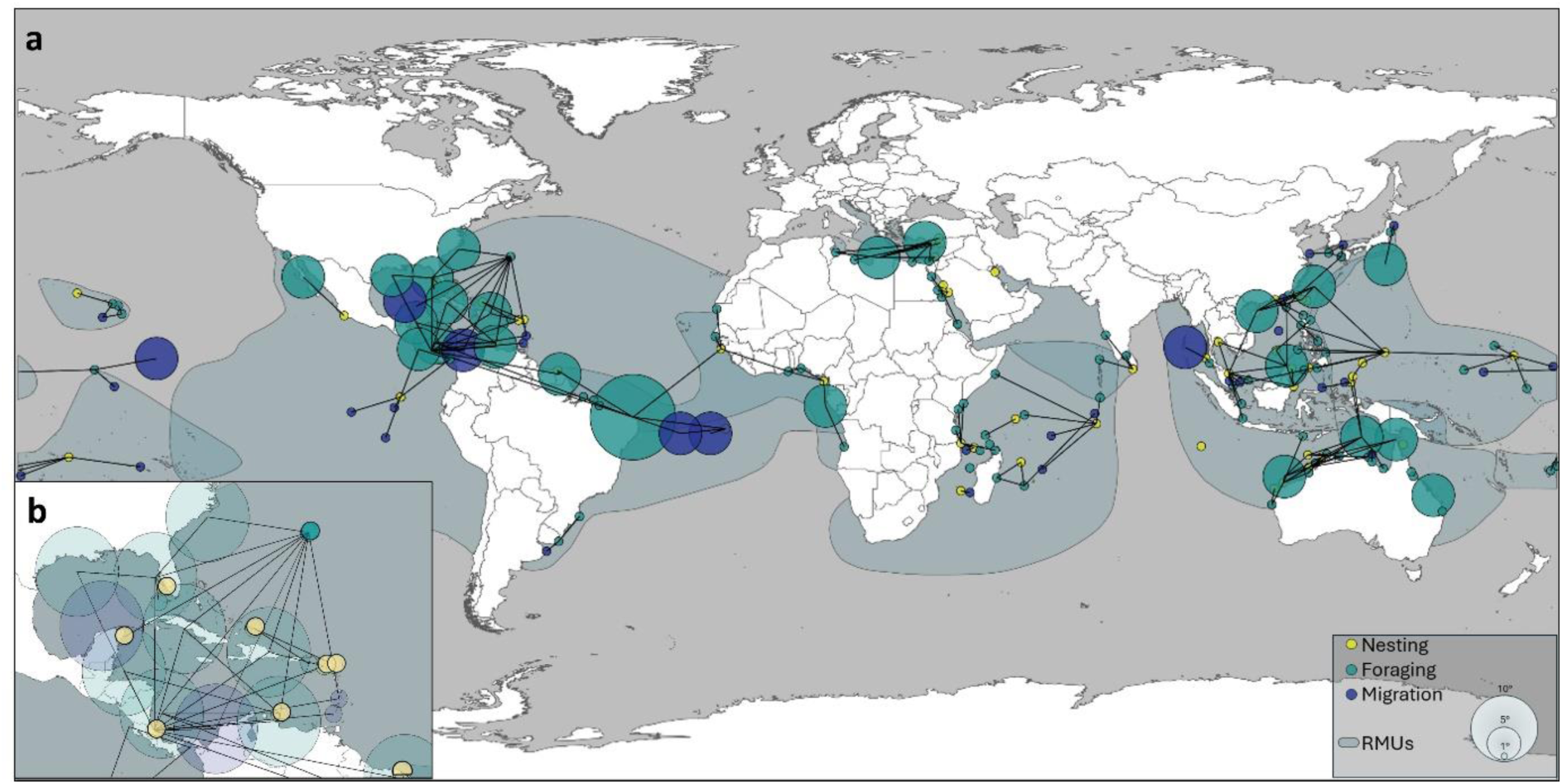
Analysis of migratory connectivity for green turtles (Chelonia mydas). a) Worldwide connectivity network based on published literature published (1990-2022) classified by behaviour (Nesting, Foraging, Migrating). b) Inset showing the connectivity network in the North Atlantic Ocean and Greater Caribbean Sea classified by behaviour. Lines between metasites denote connections based on the tracking or recapture of tagged turtles moving between sites. Blue shaded areas represent Regional Management Units (RMUs; Wallace et al. 2023).

We identified five nesting sites that showed no connections to any other site, such as Qaru Island in Kuwait and the Cocos (Keeling) Islands in the Indian Ocean, that showed no connections to any other site. This implies local connectivity where turtles did not migrate far (∼100 km), establishing residency within the area where they were initially tracked. The remaining nesting sites had connections spanning from 200 km to almost 6,000 km (Figure 4C), which included between 1 and 13 direct connections with other sites.

**Figure 4.**
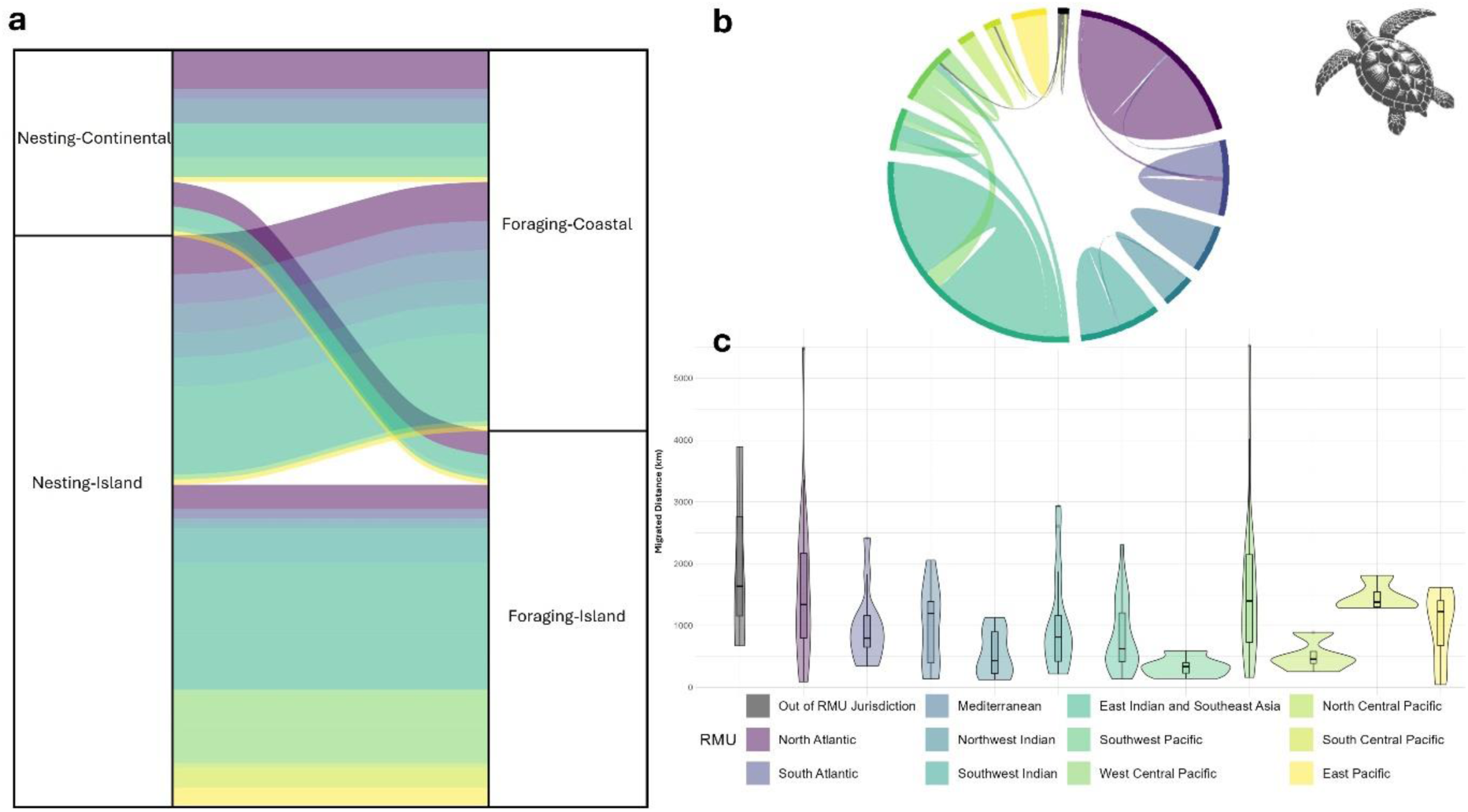
Green turtles (*Chelonia mydas*) migratory and connectivity movement description, based on satellite telemetry. a) Connectivity between different types of nesting and foraging habitats by Regional Management Unit (RMU). b) Connectivity and movement of green turtles within and between RMUs. c) Distance travelled by green turtles evaluated by RMU.

Tracked green turtles nested on sandy beaches, predominantly on small islands (87.1%), and continental beaches (12.9%). Although, only a few continental rookeries were reported in the Indian (11.1%) and Pacific (14.3%) oceans, almost half of the studied nesting sites in the Atlantic Ocean (46.7%) were located on continental beaches. Migratory movements differed among regions depending on the geomorphological characteristics of nesting sites (Figure 4a). In the Caribbean Sea and North Atlantic (RMU 1), connections primarily linked continental nesting sites on the mainland to foraging sites in coastal areas dispersed throughout the region, though some turtles migrated towards foraging grounds located around islands and atolls (Figure 4a). Conversely, connectivity patterns in the Indian and Pacific basins (RMUs 4 to 11) showed a combination of strategies used by green turtles in these areas. Most of the island-nesting turtles migrated towards shallow waters surrounding other islands, whereas a smaller proportion of nesting turtles travelled to seagrass meadows in coastal continental areas (Figure 3a, 4a). Migratory strategies within the Mediterranean Sea (RMU 3) revealed all turtles moving to coastal foraging areas, primarily offshore African nations (Figure 3a).

### Migratory connectivity and Regional Management Units

Movements recorded for green turtles were almost entirely within RMU polygons. Research has been conducted in every RMU, and most monitored turtles were initially tagged and released within the boundaries of RMUs. Though most metasites and connections remained within unique RMUs, we also found nesting and foraging sites located in regions where different RMUs overlapped. Thus, connecting routes from these sites described movement from turtles either crossing RMUs or moving outside of these areas (Figure 3a). Further, we identified 13.7% of connections between key nesting and foraging sites crossed RMU borders (Figure 4b). In northern Australia, we found green turtles moving from nesting sites to foraging areas, crossing from the Southwest Pacific to the West Central Pacific RMUs.

We registered similar movements from a few nesting populations crossing RMU boundaries in every ocean basin except the Red Sea (Northwest Indian, RMU 4), the Hawaiian archipelago (North Central Pacific, RMU 9), and the Mediterranean Sea (RMU 3), which had relatively short migrations that remained within their respective RMUs (Figure 4b, 4c).

## Discussion

Green turtles are the most studied species of marine turtles and the most data rich marine migratory species (Kot et al. 2022, Bentley et al. 2025). Driven by increasingly affordable and effective telemetry technology, the number of studies addressing migratory behaviour of green turtles has increased over the past three decades (Godley et al. 2008; Jeffers & Godley 2016). We synthetised this information and developed a novel network model that provides ecological insights into regional differences in life histories, identifies geographic and demographic gaps in sampling, and shows baseline information on connectivity. We identified a novel typology of life history movement patterns, presenting two distinct nesting site selection strategies, with opposite continent-island (or island-continent) migrations, connecting rookeries and foraging grounds. We also found large variations in the distance between nesting and foraging areas. Most connections showed movement traversing national jurisdictions, with individual green turtle rookeries connecting ecosystems in up to 13 countries. Further, long-distance migrations for some populations included occasional movement across different Regional Management Units.

### Global migratory connectivity

Historically, the ecology of green turtles has been modelled to assess abundance and distribution of populations at nesting beaches (e.g. Mazaris et al. 2017; Eckert & Eckert 2019, Kot et al. 2023). These nesting beaches are strategic sites acting as key nodes for the migration of green turtles, not only as a point of origin for hatchling dispersal (Scott et al. 2014; Le Gouvello et al. 2020), but as source areas to track breeding migrations (Whiting et al. 2008). In this connectivity network, we preserve nesting rookeries as unique sites and show migration pathways for different populations worldwide. Connections from nesting sites ranged from resident turtles remaining in the area where they were encountered to routes linking rookeries with various and distant foraging locations. Indeed, connections between nesting and foraging sites for green turtles have been reported up to 10,000 km apart (Shimada et al. 2019), nonetheless, we found that most post-nesting migrations range from a few hundred to almost six thousand kilometres (Figure 4c). This pattern suggests that most nesting aggregations of green turtles are being supported by multiple, dispersed foraging areas (Figure 3a). Further, long-term monitored nesting populations have shown connections to dispersed foraging grounds over several decades in different regions globally (Troëng et al. 2005, Stokes et al. 2015, Ferreira et al. 2021). Predominantly herbivorous (Esteban et al. 2020), green turtles are highly dependent on seagrass meadows and thus declines in seagrass abundance and shrinking of distributions can cause changes in the demographics and ecology of green turtle populations (Meylan et al. 2022). Therefore, nesting aggregations dispersing over several foraging areas constitute a plausible portfolio strategy to improve their chances of survival by offsetting the risk of an individual seagrass meadow being depleted (Fourqurean et al. 2019).

Conversely, the aggregation of foraging sites into metasites has revealed key areas where turtles from different rookeries migrate to forage. Foraging aggregations of green turtles are commonly composed of a combination of genetically distinct stocks (Jensen et al. 2019; van der Zee et al. 2019). The mixing of genetic stocks in foraging grounds could be interpreted as the result of a combination of effects of oceanic currents on the dispersal of hatchlings, together with the size and proximity of the rookeries from where the turtles originally migrated (Luke et al. 2004). Aggregated 5° foraging metasites in this network are highly interconnected with several nesting beaches and other metasites (Figure 3a), and we found individual foraging sites connected to up to nine different rookeries. For instance, in the Pacific Basin, metasites off the northern coast of Borneo act as connectivity nodes, linking five rookeries with nearby foraging metasites. Similarly, foraging metasites on the north coast of Australia are connected to nesting and foraging sites in Western Australia, Indonesia and the Philippines. Such metasites constitute small, complex networks, each connecting diverse nesting sites and foraging locations across several national jurisdictions and even crossing RMUs’ limits (Figure 4b), expanding their boundaries to overlapping areas.

### Connectivity strategies

Green turtles tend to nest on sandy, tropical beaches either on continental shores or islands (Seminoff et al. 2015; Figure 2c). Our network model revealed a novel typology of the life history for green turtles, highlighting differences in the selection of distinctive nesting habitats across populations, and their connectivity to defined foraging areas either in coastal ecosystems or insular areas in shallow water near atolls, sand banks or small islands.

Though most of the nesting sites for green turtles were located on islands, we found foraging sites to be equally distributed in continental coastal areas and insular waters (Figure 4a).

Identifying these areas is key to assess the population-level impacts of threats (Hays et al. 2025, Wallace et al. 2025), and it has important implications for the adaptability of green turtles to future threats. The loss of nesting sites (McClenachan et al. 2006) and environmental factors have a permanent, irreversible effect on sea turtle populations by limiting their reproductive output (Saragoça et al. 2020). Similarly, the continued decline of seagrass meadows (Orth et al. 2006; Waycott et al. 2009), has a great impact on developing and foraging areas, hindering green turtles’ capacity to overcome emerging threats on food resources and habitat alterations, influencing their adaptation ability and survival.

We identified key nodes of connectivity comprising nesting sites in Australia, the South Atlantic, Hawaii, Micronesia, and the Indian Ocean, among others, where green turtles’ rookeries were primarily located on exposed sandy beaches. Island and atoll rookeries are characterised by reduced vegetation cover and modest topographic profiles. These parameters directly affect conditions during the incubation period, increasing risk of inundation due to elevated groundwater levels, influencing survival rates, increasing temperature profiles, affecting hatching success, and the sex ratios on these beaches (Conrad et al. 2011; Howard et al. 2014; Hays et al. 2017; Hernández-Cortés et al. 2018; Monsinjon et al. 2022). Though we found a high degree of connectivity amongst insular rookeries, as well as diverse foraging sites, the challenge to adapt to projected changes in temperature for green turtles nesting on islands is of concern for several populations (Fuentes et al. 2024).

In contrast, connectivity of green turtles across the Caribbean Sea presented a different dynamic. Although some of the monitored nesting sites for green turtles are located on islands across the Antilles, most of the larger nesting sites (by several orders of magnitude) in this region are located on coastal beaches on the continental shore of Costa Rica, Florida and Mexico (Figure 3b) (Broderick et al. 2006). While green turtle populations across the globe are susceptible to impacts of global warming and sea level rise (Fuentes et al. 2010; Martins et al. 2022), the availability of bigger and naturally shaded beaches in continental sites may provide enough protection for green turtles to mitigate these threats (Fuentes et al. 2020; Reboul et al. 2021). Unfortunately, climate change impacts are not the only threat for green turtles in the region. Effects of ongoing anthropogenic activities are currently the major pressure on green turtles in the Greater Caribbean (Senko et al. 2022), with egg harvesting and marine turtle meat consumption still widespread in local communities and traditionally used by Indigenous and Afro-Caribbean communities (Meylan et al. 2013, Barrios-Garrido et al. 2017, Lagueux et al. 2017).

Within our network, the Caribbean Sea is the most data-rich area for migratory connectivity of green turtles. We identified two primary nodes of connectivity: Tortuguero Beach in Costa Rica, which is the largest green turtle rookery in the western hemisphere (Restrepo et al. 2023); and a developmental area in shallow waters on the Bermuda Platform, which is >1,000 km from other foraging grounds, and where no adult green turtles are found (Meylan et al. 2022). Although green turtles have not been detected migrating directly between Tortuguero and Bermuda, connections from each of these sites lead to similar areas (Troëng et al. 2005; Meylan et al. 2011) where mature turtles and large subadults cohabit the same habitats, in preparation for future reproductive migrations (Shimada et al. 2016). The Miskito Cays in Nicaragua, the southern coast of Cuba, and waters around the Yucatan Peninsula in Mexico are some of the largest aggregated foraging metasites we found in the Caribbean.

Seagrass meadows in these locations are the primary food source for adult green turtles in the region (Moncada et al. 2006; Lagueux et al. 2017), supporting some of the largest nesting populations worldwide.

### Migratory connectivity and Regional Management Units

RMUs provide a framework to define subpopulations of marine turtles within their near-global distribution (Wallace et al. 2010). Our connectivity network shows migratory behaviour that is broadly aligned with defined RMUs. This is not surprising, given that RMU boundaries and our network model were both established using some of the same literature (Wallace et al. 2023). Movement pathways and routes synthesised in the current study reveal high connectivity among distant key sites for green turtles around the world. For instance, nesting turtles in Micronesia in the Pacific Ocean travelled >5,000 km, crossing Filipino and Vietnamese waters to reach seagrass meadows off the coast of Johor province in Southeast peninsular Malaysia. Similarly, green turtle migratory patterns described in this region highlight the movement of nesting turtles in rookeries located in the West Central Pacific RMU, towards foraging habitats within the Southwest Pacific, the East Indian and Southeast Asia RMUs (Figure 3a). Connections crossing RMU boundaries are also shown in overlapping areas of RMUs in the Atlantic and Indian Ocean (Figure 4b), demonstrating that green turtles travel long distances and link key habitats across different RMUs.

A key goal of RMUs is to focus management efforts in specific areas where populations of marine turtles reside (Wallace et al. 2023). We show connections linking nesting sites and foraging areas within different RMUs. Similarly, we found green turtle movements compiled in this study showed migratory pathways of breeding individuals crossing several national boundaries and occasionally entering areas beyond national jurisdictions (except for turtles tagged in Australia, and a few foraging grounds on the coast of Brazil). In contrast to leatherback turtles that embark on seasonal trans-oceanic reproductive migrations and spend >78% of their time in the high seas (Harrison et al. 2018), green turtles generally prefer coastal waters, though few exceptions have been reported for populations in all oceanic basins. This constrains their movement, limiting their time in areas beyond national jurisdiction to a few weeks each time they embark on a breeding migration. These movements reflect the complexity of marine turtle migratory connectivity and suggest that their management might benefit from knowledge synthesis and collaboration over multiple RMUs (Mazaris et al. 2017).

### Gaps and limitations

Tracking green turtle movements in the vastness of the ocean presents a series of challenges for researchers, which should be considered when interpreting this global connectivity network. Lack of information on movement for juveniles and male turtles means there is a bias towards reproductive females (Kot et al. 2023). Although the proportion of different classes has increased in recent studies compared to previous reviews (Godley et al. 2008; Rees et al. 2016), nesting females are still the most sampled group (70.1%) in movement or connectivity studies. This bias likely leads to under representation of key habitats and migratory routes for entire populations. Further, the number of studies assessing movement of green turtles covers ∼10% of populations worldwide. In our analysis we have included all the migratory information available in the scientific literature published in English over the past 33 years, including post-nesting migrations from green turtles departing 124 nesting sites. Nonetheless, >1,100 nesting sites have been reported for green turtles (Pritchard. 2011, Kot et al. 2023). Finally, due to limitations of current tag batteries and the extensive nature of the migration patterns for most green turtles, it is likely we have not captured the full migratory route of many tagged turtles. Thus, when describing the green turtle global connectivity network, we considered broad-scale connections from identified nodes, weaving the minimum known scope of connectivity for green turtles at a global scale.

These limitations are a present throwback not just to study marine turtles but all marine migratory species moving between breeding and foraging delimited areas. Connectivity models presented by kot et al. (2023) and Bentley et al. (2025), as well as the present network, showcase the potential to use existing information at a relative low cost, to effectively inform conservation measures at a regional, and global scale incorporating key habitat for vulnerable and highly migratory species.

## Conclusion

We compiled the largest available dataset describing movement of green turtles worldwide and present the first global model of their migratory connectivity. We uncovered key knowledge gaps on movement patterns of multiple life stages, and highlighted rookeries and foraging grounds as key connectivity nodes for green turtle populations. We aggregated foraging sites to better understand connections among these habitats and to better inform local and regional management for conservation of marine areas globally. Our network model revealed a novel typology of nesting green turtles which will have serious implications for the species facing adaptation to climate change threats and which should be further investigated to understand their implications for conservation.

## Data Accessibility

Raw literature review datasheets, datafiles and scripts are available for review at GITHUB, an archived version of these data will be available in Zenodo for the published version of this article. https://github.com/jaime-restrepo/Cm_Connectivity_Network.git

## Notes

https://github.com/jaime-restrepo/Cm_Connectivity_Network.git

### Competing Interest Statement

The authors have declared no competing interest.

### Funding

- German Federal Ministry for the Environment, Nature Conservation, Nuclear Safety and Consumer Protection: DP240101026
- Global Ocean Biodiversity Initiative grant from the International Climate Initiative: 16_IV_049_Global_A

