## Supplemental Data 1 for "Ecological insights and management implications of the global migratory connectivity of green turtles"

***Supplementary Material***

***Table S1.*** *Information on movement behaviour, tracking technique, sex, and life-stage classes of green turtles from published literature (1990–2022).*

| Movement Behaviour | Tracking Technique | Tags Applied | Sex | | | Life Stage | | | | | References |
|---|---|---|---|---|---|---|---|---|---|---|---|
| | | | Female | Male | Und. | Adult | Adult Breeding | Subadult | Juvenil | Unknown | |
| Post-Nesting Migration | Mark-Recapture | 5077 | 5076 | - | 1 | - | 5074 | 3 | - | - | 4, 27, 29, 42, 54, 62, 68, 75, 112 |
| | Satellite Telemetry | 404 | 404 | - | - | 2 | 375 | 27 | - | - | 5, 7, 11, 12, 18, 21, 23, 28, 30, 40, 43, 45, 57, 60, 73, 76, 77, 78, 79, 81, 86, 87, 90, 91, 93, 94, 95, 96, 97, 100, 103 |
| | Dive Loggers | 10 | 10 | - | - | - | 10 | - | - | - | 88 |
| Habitat Use | Mark-Recapture | 555 | 32 | 45 | 478 | 123 | - | 193 | 239 | - | 84 |
| | Satellite Telemetry | 192 | 113 | 19 | 60 | 84 | 27 | 0 | 49 | 32 | 1, 9, 25, 32, 35, 58, 71, 83, 98, 102, 104, 105 |
| | Acoustic Telemetry | 86 | 23 | 25 | 38 | 16 | 42 | 13 | 15 | - | 46, 48, 61, 64, 67, 99, 72 |
| Foraging | Mark-Recapture | 277 | 170 | 31 | 76 | - | 140 | 34 | 41 | 62 | 65, 66, 85,101 |
| | Satellite Telemetry | 103 | 64 | 2 | 37 | - | 66 | - | 30 | 7 | 2, 16, 17, 52, 56 |
| | Acoustic Telemetry | 93 | 19 | 16 | 58 | | 1 | 22 | 34 | 36 | 33, 80, 107 |
| | Dive Loggers | 1 | 1 | - | - | - | 1 | - | - | - | 50 |
| Swimming Behaviour / Migratory Movements | Mark-Recapture | 246 | 187 | - | 59 | 2 | 187 | - | 4 | 53 | 10, 15, 108 |
| | Satellite Telemetry | 286 | 175 | 24 | 87 | 42 | 120 | 20 | 87 | 17 | 6, 8, 13, 19, 22, 24, 31, 36, 37, 38, 41, 44, 47, 50, 53, 55, 59, 69, 70, 82, 89, 92, 104, 109, 110, 111, 113 |
| | Dive Loggers | 4 | 4 | - | - | - | 4 | - | - | - | 110 |
| Interactions / Fisheries | Satellite Telemetry | 7 | - | - | 7 | - | - | - | 7 | - | 20 |
| | Combined Techniques | 10 | 10 | - | - | - | - | - | 10 | - | 74 |
| Experimental Trials | Satellite Telemetry | 6 | 4 | - | 2 | - | - | - | - | - | 14, 34, 39 |
| | Combined Techniques | 3 | - | - | 3 | - | - | - | - | 3 | 3 |

### *Connectivity network references*

56. Kennett, R., Munungurritj, N., & Yunupingu, D. (2004). Migration patterns of marine turtles in the Gulf of Carpentaria, northern Australia: implications for Aboriginal management. *Wildlife Research*, *31*(3), 241-248. https://doi.org/10.1071/WR03002
57. Kittiwattanawong, K., Chantrapornsyl, S., Sakamoto, W., & Arai, N. (2002). Tracking of green turtles Chelonia mydas in the Andaman Sea using platform transmitter terminals. Phuket Marine Biological Center Research Bulletin (Thailand), (64).
58. Klain, S., Eberdong, J., Kitalong, A., Yalap, Y., Matthews, E., Eledui, A., ... & Kemesong, P. (2007). Linking Micronesia and Southeast Asia: Palau sea turtle satellite tracking and flipper tag returns. *Marine Turtle Newsletter*, *118*, 9-11.
59. Kolinski, S. P., Cruce, J., Parker, D. M., Balazs, G. H., & Clarke, R. (2014). Migrations and conservation implications of post-nesting green turtles from Gielop Island, Ulithi Atoll, Federated States of Micronesia. *Micronesica*, *4*, 1-9.
60. Lamont, M. M., Fujisaki, I., Stephens, B. S., & Hackett, C. (2015). Home range and habitat use of juvenile green turtles (Chelonia mydas) in the northern Gulf of Mexico. *Animal Biotelemetry*, *3*, 1-12. https://doi.org/10.1186/s40317-015-0089-9
61. Levy, Y., Keren, T., Leader, N., Weil, G., Tchernov, D., Rilov, G., (2017). Spatiotemporal hotspots of habitat use by loggerhead (Caretta caretta) and green (Chelonia mydas) sea turtles in the Levant basin as tools for conservation. Marine Ecology Progress Series 575, 165–179.. https://doi.org/10.3354/meps12146
62. Liew, H. C., Chan, E. H., Luschi, P., Papi, F., & Papi, F. (1995). Satellite tracking data on Malaysian green turtle migration. *Rendiconti Lincei. Scienze Fisiche e Naturali*, *6*(3), 239-246. https://ui.adsabs.harvard.edu/link_gateway/1995RLSFN...6..239L/doi:10.1007/BF03001671
63. Lima, E. H. S. M., Lagueux, C. J., Castro, D., & Marcovaldi, M. A. (1999). From one feeding ground to another: Green turtle migration between Brazil and Nicaragua. *Marine Turtle Newsletter*, *85*(10).
64. Luschi, P., Hays, G. C., Del Seppia, C., Marsh, R., & Papi, F. (1998). The navigational feats of green sea turtles migrating from Ascension Island investigated by satellite telemetry. *Proceedings of the Royal Society of London. Series B: Biological Sciences*, *265*(1412), 2279-2284. https://doi.org/10.1098/rspb.1998.0571
65. MacDonald, B. D., Madrak, S. V., Lewison, R. L., Seminoff, J. A., & Eguchi, T. (2013). Fine scale diel movement of the east Pacific green turtle, Chelonia mydas, in a highly urbanized foraging environment. *Journal of Experimental Marine Biology and Ecology*, *443*, 56-64. https://doi.org/10.1016/j.jembe.2013.02.033
66. Madrak, S. V., Lewison, R. L., Seminoff, J. A., & Eguchi, T. (2016). Characterizing response of East Pacific green turtles to changing temperatures: using acoustic telemetry in a highly urbanized environment. *Animal Biotelemetry*, *4*, 1-10. https://doi.org/10.1186/s40317-016-0114-7
67. Mansfield, K. L., Wyneken, J., & Luo, J. (2021). First Atlantic satellite tracks of 'lost years' green turtles support the importance of the Sargasso Sea as a sea turtle nursery. *Proceedings of the Royal Society B*, *288*(1950), 20210057. https://doi.org/10.1098/rspb.2021.0057
68. McClellan, C. M., Read, A. J., Price, B. A., Cluse, W. M., & Godfrey, M. H. (2009). Using telemetry to mitigate the bycatch of long-lived marine vertebrates. *Ecological Applications*, *19*(6), 1660-1671. https://doi.org/10.1890/08-1091.1
69. McClellan, C. M., & Read, A. J. (2009). Confronting the gauntlet: understanding incidental capture of green turtles through fine-scale movement studies. *Endangered Species Research*, *10*, 165-179. https://doi.org/10.3354/esr00199

98. Snape, R. T., Beton, D., Davey, S., Godley, B. J., Haywood, J., Omeyer, L. C., Ozkan, M., & Broderick, A. C. (2022). Mediterranean green turtle population recovery increasingly depends on Lake Bardawil, Egypt. Global Ecology and Conservation, 40, e02336. https://doi.org/10.1016/j.gecco.2022.e02336
99. Snoddy, J. E., & Williard, A. S. (2010). Movements and post-release mortality of juvenile sea turtles released from gillnets in the lower Cape Fear River, North Carolina, USA. Endangered Species Research, 12(3), 235-247. https://doi.org/10.3354/esr00305
100. Stokes, K. L., Broderick, A. C., Canbolat, A. F., Candan, O., Fuller, W. J., Glen, F., Levy, Y., Rees, A. F., Rivol, G., Snape, R. T., Tchernov, D., & Godley, B. J. (2015). Migratory corridors and foraging hotspots: critical habitats identified for Mediterranean green turtles. *Diversity and Distributions*, *21*(6), 665-674. https://doi.org/10.1111/ddi.12317
101. Stringell, T. B., Clerveaux, W. V., Godley, B. J., Phillips, Q., Ranger, S., Richardson, P. B., ... & Broderick, A. C. (2015). Protecting the breeders: research informs legislative change in a marine turtle fishery. *Biodiversity and conservation*, *24*, 1775-1796. https://doi.org/10.1007/s10531-015-0900-1
102. Swimmer, Y., Arauz, R., McCracken, M., McNaughton, L., Ballestero, J., Musyl, M., Bigelow, K., & Brill, R. (2006). Diving behavior and delayed mortality of olive ridley sea turtles Lepidochelys olivacea after their release from longline fishing gear. Marine Ecology Progress Series, 323, 253-261. https://doi.org/10.3354/meps323253
103. Troëng, S., Evans, D. R., Harrison, E., & Lagueux, C. J. (2005). Migration of green turtles Chelonia mydas from Tortuguero, Costa Rica. Marine Biology, 148(2), 435-447. https://doi.org/10.1007/s00227-005-0076-4
104. Türkecan, O., & Yerli, S. V. (2011). Satellite tracking of adult green sea turtles from Turkey: A long distance diary. *Marine Turtle Newsletter*, (131), 38.
105. van de Merwe, J. P., Ibrahim, K., Lee, S. Y., & Whittier, J. M. (2009). Habitat use by green turtles (Chelonia mydas) nesting in Peninsular Malaysia: local and regional conservation implications. *Wildlife Research*, *36*(7), 637-645. https://doi.org/10.1071/WR09099
106. Vélez-Rubio, G. M., Cardona, L., López-Mendilaharsu, M., Souza, G. M., Carranza, A., Campos, P., González-Paredes D., & Tomás, J. (2018). Pre and post-settlement movements of juvenile green turtles in the Southwestern Atlantic Ocean. *Journal of Experimental Marine Biology and Ecology*, *501*, 36-45. https://doi.org/10.1016/j.jembe.2018.01.001
107. Waayers, D. A., Smith, L. M., & Malseed, B. E. (2011). Inter-nesting distribution of green turtles (Chelonia mydas) and flatback turtles (Natator depressus) at the Lacepede Islands, Western Australia. *Journal of the Royal Society of Western Australia*, *94*(2), 359.
108. Weber, N., Weber, S. B., Godley, B. J., Ellick, J., Witt, M., & Broderick, A. C. (2013). Telemetry as a tool for improving estimates of marine turtle abundance. *Biological Conservation*, *167*, 90-96. https://doi.org/10.1016/j.biocon.2013.07.030
109. Webster, E. G., Hamann, M., Shimada, T., Limpus, C., & Duce, S. (2022). Space-use patterns of green turtles in industrial coastal foraging habitat: Challenges and opportunities for informing management with a large satellite tracking dataset. *Aquatic Conservation: Marine and Freshwater Ecosystems*, *32*(6), 1041-1056. https://doi.org/10.1002/aqc.4239
110. Whiting, S. D., & Miller, J. D. (1998). Short term foraging ranges of adult green turtles (Chelonia mydas). *Journal of Herpetology*, 330-337. https://www.jstor.org/stable/1565446
111. Whiting, S. D., Murray, W., Macrae, I., Thorn, R., Chongkin, M., & Koch, A. U. (2008). Non-migratory breeding by isolated green sea turtles (Chelonia mydas) in the Indian Ocean: biological and conservation implications. *Naturwissenschaften*, *95*(4), 355-360. https://doi.org/10.1007/s00114-007-0327-y

112. Yasuda, T., & Arai, N. (2005). Fine-scale tracking of marine turtles using GPS-Argos PTTs. *Zoological Science*, *22*(5), 547-553. https://doi.org/10.2108/zsj.22.547

113. Yasuda, T., Tanaka, H., Kittiwattanawong, K., Mitamura, H., Klom-in, W., & Arai, N. (2006). Do female green turtles (Chelonia mydas) exhibit reproductive seasonality in a year-round nesting rookery?. *Journal of Zoology*, *269*(4), 451-457. https://doi.org/10.1111/j.1469-7998.2006.00134.x
